# Circuit models of low dimensional shared variability in cortical networks

**DOI:** 10.1101/217976

**Authors:** Chengcheng Huang, Douglas A. Ruff, Ryan Pyle, Robert Rosenbaum, Marlene R. Cohen, Brent Doiron

## Abstract

Trial-to-trial variability is a reflection of the circuitry and cellular physiology that makeup a neuronal network. A pervasive yet puzzling feature of cortical circuits is that despite their complex wiring, population-wide shared spiking variability is low dimensional with all neurons fluctuating en masse. Previous model cortical networks are at loss to explain this global variability, and rather assume it is from external sources. We show that if the spatial and temporal scales of inhibitory coupling match known physiology, model spiking neurons internally generate low dimensional shared variability that captures the properties of *in vivo* population recordings along the visual pathway. Shifting spatial attention into the receptive field of visual neurons has been shown to reduce low dimensional shared variability within a brain area, yet increase the variability shared between areas. A top-down modulation of inhibitory neurons in our network provides a parsimonious mechanism for this attentional modulation, providing support for our theory of cortical variability. Our work provides a critical and previously missing mechanistic link between observed cortical circuit structure and realistic population-wide shared neuronal variability and its modulation.

## Introduction

The trial-to-trial variability of neuronal responses gives a critical window into how the circuit structure connecting neurons drives brain activity^1^. This idea combined with the widespread use of population recordings has prompted deep interest in how variability is distributed over a population^2,3^. There has been a proliferation of data sets where the shared variability over a population is low dimensional^4–9^, meaning that neuronal activity waxes and wanes as a group. In accord, one dimensional measures such as local field potentials^10,11^ and summed population firing rates can predict a majority of pairwise correlations^9,12^. Further, the synthesis of diverse population datasets paints a picture where low dimensional shared variability is a signature of cognitive state, such as overall arousal, task engagement and attention^2,3,13^, as well as predictive of behavioral performance^14^. Such low dimensional dynamics portend a theory for the genesis and modulation of shared population variability in recurrent cortical networks.

Theories of cortical variability can be broadly separated into two categories: ones where variability is internally generated through recurrent network interactions (Fig. 1a, left) and ones where variability originates external to the network (Fig. 1a, middle). Networks of spiking neuron models where strong excitation is balanced by opposing recurrent inhibition produce high single neuron variability through internal mechanisms^15–17^. However, these networks famously enforce an asynchronous state, and as such fail to explain population-wide shared variability^18^. This lack of success is contrasted with the ease of producing arbitrary correlation structure from external sources. Indeed, many past cortical models assume a global fluctuation from an external source^3,7,19–21^, and accurately capture the structure of population data. However, such phenomenological models are circular, with an assumption of variability from an unobserved source explaining the variability in a recorded population. Thus, while neuronal variability has a rich history of study, there remains an empoverished mechanistic understanding of the low dimensional structure of population-wide variability^22^.

**Figure 1:**
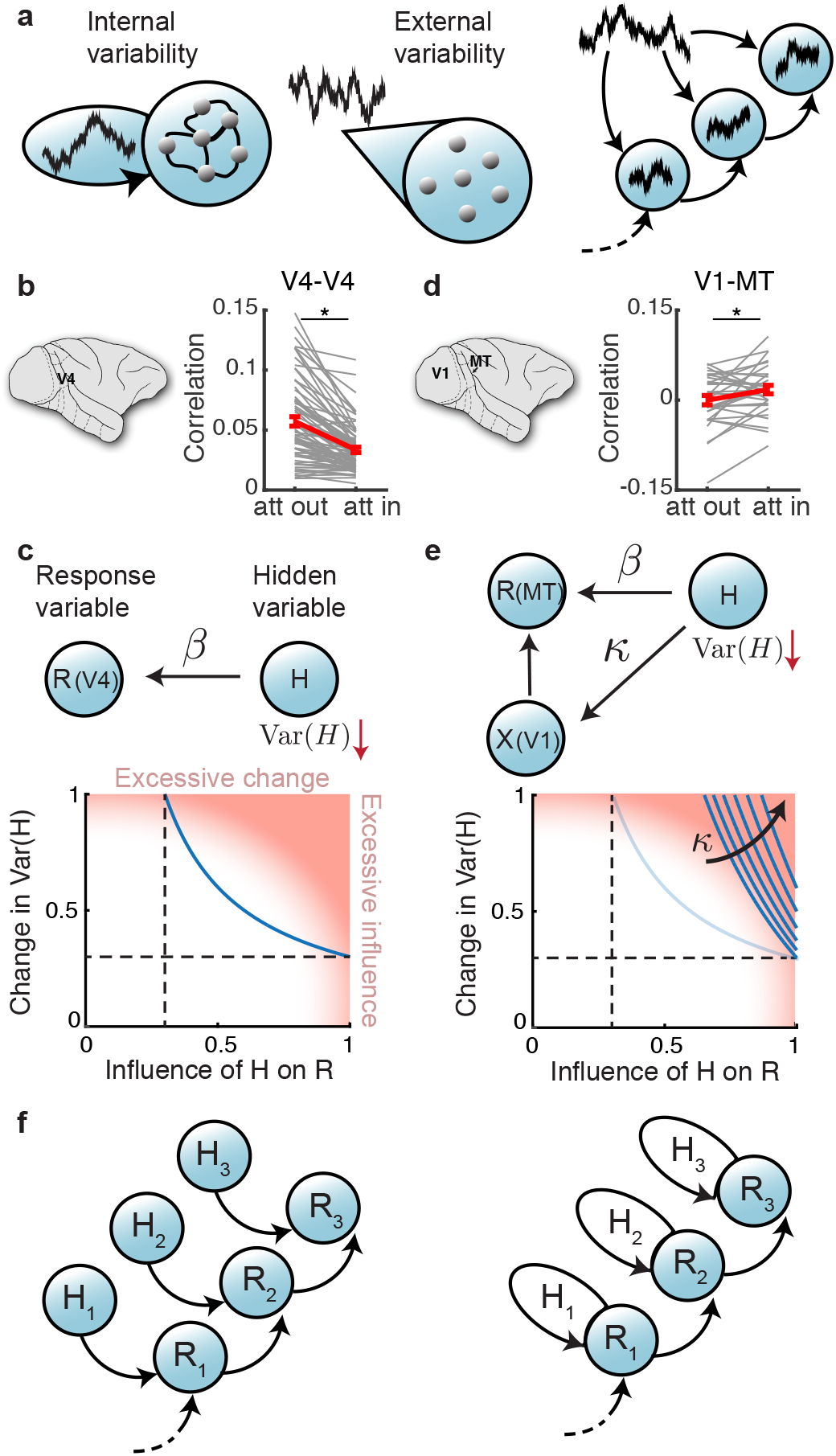
Models of shared variability. **a**, Variability may either be internally generated within a population (left) or externally imposed upon a population (middle). New model constraints emerge by accounting how variability is distributed and modulated across several populations (right). **b**, Mean spike count correlation *r_SC_* per session obtained from multi-electrode array recording from V4 was smaller when attention was directed into the receptive fields of recorded neurons (n=74 sessions, two-sided Wilcoxon rank-sum test between attentional states *P* = 3.3 × 10^−6^, reproduced from^24^). Grey lines are individual session comparisons and the red line is the mean comparison across all sessions (error bars represent the SEM). **c**, Top: hidden variable model where the response variability *R* (modeling V4) comes from a hidden variable *H* with influence *β*. Bottom: the attention-mediated reduction in *r_SC_* gives a constraint that is a trade-off between the reduction in Var(*H*) and *β* (blue curve). **d**, Same as **b** for the mean spike count correlation *r_SC_* between V1 units and MT units per session (n=32 sessions, paired-sample *t*-test *P* = 0.0222; data reproduced from^23^). **e**, Top: hidden variable model for connected areas *X* (modeling V1) and *R* (modeling MT); *H* projects to *X* with strength *κ*. Bottom: the attention mediated changes in *r_SC_* give further constraints on *H* with the increase in *κ* indicated. Light blue curve is the same as that in **c** for comparison. **f**, Schematic for external (left) and internal (right) models of shared variability (H) along a processing hierarchy (R).

Determining whether output variability is internally generated through network interactions or externally imposed upon a network is a difficult problem, where single area population recordings may preclude any definitive solution (Fig. 1a, left vs middle). In this study we consider attention-mediated shifts in population variability obtained from simultaneous recordings of neuron pairs both within and between visual areas^23,24^. Attention reduces within area correlations (area V4) while simultaneously increasing between area correlations (areas V1 and MT), thereby providing a novel constraint for how shared variability is distributed within and between neuronal populations (Fig. 1a, right). We present analysis showing that such a differential correlation modulation is difficult constraint to satisfy with a model where fluctuations are strictly external to the network. We thus focus our modeling on networks where population-wide shared variability can be internally generated.

The asynchronous solution of classical balanced networks necessitates that inhibition dynamically tracks and cancels any correlations steaming from recurrent excitation^18^. This requirement has forced theorists to assume that the timecourse of inhibitory synapses is faster than that of excitatory synapses^16,18,25–27^, at odds with recorded synaptic physiology^28^. Recently, we have extended the theory of balanced networks to include a spatial component to network architecture^25,26,29^ and found network solutions where firing rate balance and asynchronous dynamics are decoupled from one another^26^. In this study, we consider multi-area models of spatially distributed balanced networks and show that when inhibition has slower kinetics than excitation these networks, matching physiology, they internally produce low dimensional population-wide variability. Unlike networks that lack spatial structure, these networks produce spiking activity that robustly captures the rich diversity of firing rate and correlated structure of real population recordings. Further, attention-mediated top-down modulation of inhibitory neurons in our model provides a parsimonious mechanism that controls population-wide variability in agreement with the within and between area experimental results.

There is a long standing research program aimed at providing a circuit-based understanding for cortical variability^1,15–17,26^. Our work is a critical advance through providing a mechanistic theory for the genesis, propagation, and modulation of realistic low dimensional population-wide shared variability based on established circuit structure and synaptic physiology.

## Results

### Externally imposed or internally generated shared variability?

Directed attention reduces the mean spike count correlation coefficient between neuron pairs in visual area V4 during an orientation detection task (Fig. 1b;^24^). In V4^5^, as with other cortices^4,6,8,9,12^, shared variability across a population is low dimensional, where coordinated fluctuations are driven by a common latent variable. Further, attention reduces pairwise correlation through attenuation of this global latent variable^5,30^. Thus motivated, we represent the aggregate population response with a scalar random variable *R* = *X* + *βH*, where *X* is a noisy stimulus input and *H* is a hidden source of fluctuations (with strength *β*; Fig. 1c, top). In this simple model the trial-to-trial fluctuations are inherited from both *X* and *H*, but we model attention as only reducing the variance of *H* (Var(*H*)). There is a large range of parameter values for our one-dimensional hidden variable model to readily explain the reduction in Var(*R*) reported in the V4 data (Fig. 1c, bottom, blue curve; see Supplemental Information). Certain parameter choices are unreasonable (pink region in Fig. 1c, bottom), such as *β* being overly large so that *R* is no longer driven by *X*, or Var(*H*) → 0 in the attended state, requiring the area that produces *H* to be silent. Fortunately, there are moderate *β* and Var(*H*) choices that capture the data (section of the blue curve that is not in the pink region in Fig. 1c, bottom). In total, a latent variable model where the variability is external to the population can account for the attentional modulation reported in our V4 data, as has been previously remarked^5,7^.

Recent multi-electrode recordings from two visual areas, MT and V1, during an attention task^23^ impose strong constraints on the simple hidden variable model. In addition to a reduction of mean spike count correlations between neuron pairs within an area (pairwise attention-related MT correlation decrease, 0.019, Wilcoxon rank sum test, *p* = 0.017; pairwise attention-related V1 correlation decrease, 0.008, Wilcoxon rank sum test, *p* = 4.9 × 10^−6^;^23^), there is an attention-mediated *increase* of spike count correlations across areas V1 and MT (Fig. 1d). Returning to the population model with *R* modeling MT, we augment the model with V1 being the input *X* = *X*_0_ + *κH* (Fig. 1e, top; see Supplementary Information). Here *κ* denotes how much the hidden variable is directly shared between areas, and *X*_0_ is the variability in *X* that is independent of *H*. The constraint curves (Fig. 1e, bottom blue) where Var(*R*) and Cov(*R, X*) match the MT-MT and V1-MT data sets require our model to assume both a large influence of *H* on *R* and a large attentional modulation of Var(*H*) (pink region in Fig. 1e, bottom). This tightening of model assumptions reflects the compromise between an attention-mediated increase in variability transfer from *X* → *R* so that Cov(*R, X*) increases and a simultaneous decrease in Var(*H*) so that Var(*R*) decreases. This compromise can be mitigated by setting *κ* to be small, meaning that a large component of the fluctuations in *R* is private from those in *X* (Fig. 1e, bottom).

While the source of private variability *H* to area *R* may still be external to the area, if we extrapolate our model to a cortical hierarchy then each area requires an external variability ‘generator’ that projects privately to that area (Fig. 1f, left). This would require a tremendous amount of neuronal hardware. A more parsimonious hypothesis is that private variability is internally generated within each area (Fig. 1f, right). Below we investigate the circuit mechanics required for low dimensional population-wide shared variability to be an emergent property within a cortical network.

### Population-wide correlations with slow inhibition in spatially ordered networks

Networks of spiking neuron models where strong excitation is balanced by opposing recurrent inhibition internally produce high single neuron variability (Fig. 2ai) with a broad distribution of firing rates (Fig. 2b, top purple curve)^16–18^. However, these networks enforce an asynchronous solution (Fig. 2c, top purple), and as such fail to explain population-wide shared variability^8,18^. Typically, balanced networks have disordered connectivity, namely where connection probability is uniform between all neuron pairs. This approximation ignores the abundant evidence that cortical connectivity is spatially ordered with a connection probability falling off with the distance between neuron pairs^31–33^. Recently we have extended the theory of balanced networks to include such spatially dependent connectivity^25,26^. Briefly, we model a two dimensional array of integrate-and-fire neurons that receive both feedforward projections from a layer of external Poisson processes and recurrent projections within the network (see Methods); connection probability of all projections decays like a Gaussian with distance. If the spatial scale of feedforward inputs is narrower than the scale of recurrent projections, the asynchronous state no longer exists^26^, giving way to a solution with spatially structured correlations (Fig. 2aii, Supplemental movie S1, Fig. S1b). Nevertheless, the mean correlation across all neuron pairs vanishes for large network size (Fig. 2c, bottom purple curve), in stark disagreement with a majority of experimental studies^2,3^ (Fig. 1b,d).

**Figure 2:**
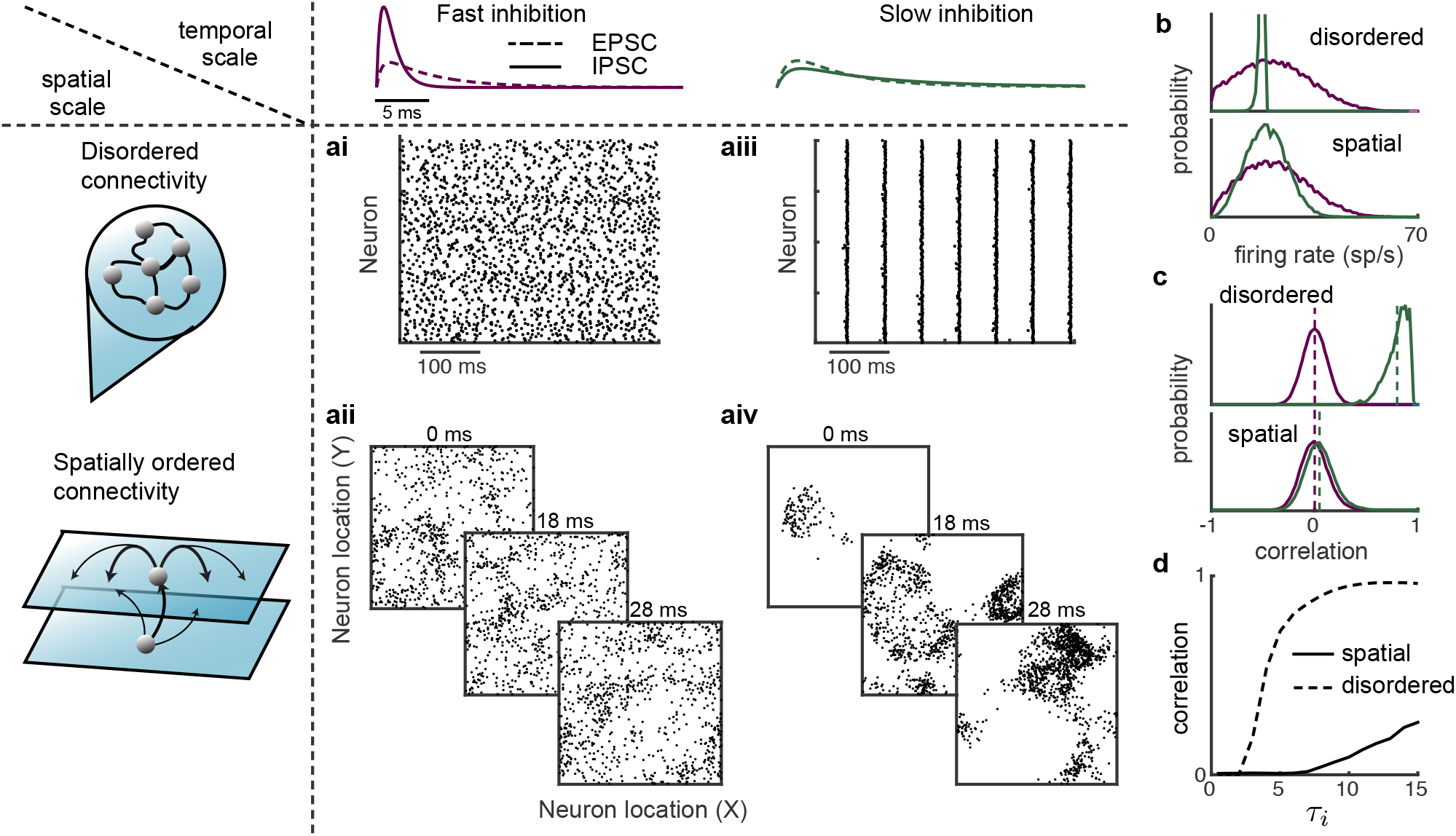
The spatial and temporal scales of synaptic coupling determine internally generated variability. **a**, Networks of excitatory and inhibitory neuron models were simulated with either disordered connectivity (ai,aiii) or spatially ordered connectivity (aii,aiv), and with either fast inhibition (*τ_i_* = 1 ms; ai,aii). or slow inhibition (*τ_i_* = 8 ms; aiii,aiv). In all models the timescale of excitation was *τ_e_* = 5 ms. In the disordered networks spike train rasters assume no particular neuron ordering. In the spatially order networks three consecutive spike raster snapshots are shown with a dot indicating that the neuron at spatial position (*x, y*) fired within one millisecond of the time stamp. **b**, Distributions of firing rates of excitatory neurons in the disordered (top) and spatially ordered (bottom) models, with faster inhibitory kinetics (purple) compared to slower inhibitory kinetics (green). **c**, Same as **b** for the distributions of pairwise correlations among the excitatory population. **d**, Mean correlation among the excitatory population as a function of the inhibitory decay time constant (*τ_i_*).

Many previous balanced network models assume that the kinetics of inhibitory conductances are *faster* than those of excitatory conductances^16–18,26,34^. However, this assumption is at odds with physiology where excitatory *α*-amino-3-hydroxy-5-methyl-4-isoxazolepropionic acid receptors (AMPA) have faster kinetics than those of the inhibitory *γ*-Aminobutyric acid receptors (GABAa)^28^. When the timescales of excitation and inhibition match experimental values in networks with disordered connectivity the activity becomes pathologic, with homogeneous firing rates (Fig. 2b, top green) and excessive synchrony (Fig. 2aiii, and c, top green). This consequence is likely the ad-hoc justification for the faster inhibitory kinetics in disordered model networks.

When a spatially ordered model has synaptic kinetics that match physiology, population-wide turbulent dynamics emerges (Fig. 2aiv, Supplemental movie S2), accompanying a small, but nonzero, mean pairwise spike count correlation across the population (*r_SC_* = 0.04). Further, firing rates are broad (Fig. 2b, bottom green curve) and pairwise correlations are reasonable in magnitude (Fig. 2c, bottom green). Indeed, as the timescale of inhibition grows, disordered networks show a rapid change in mean pairwise correlation while two dimensional spatially ordered networks show a much more gradual rise in correlation (Fig. 2d). We remark that networks constrained to one spatial dimension also produce excessive synchrony (Fig. S2), meaning that two (or more) spatial dimensions are required for robustly low but nonzero correlations. In sum, when realistic spatial synaptic connectivity is paired with realistic temporal synaptic kinetics in balanced networks, internally generated population dynamics produces spiking dynamics whose marginal and pairwise variability conform to experimental results.

### Attentional modulation of low dimensional population-wide variability

We model the V1 and MT network by extending the spatially ordered balanced networks with slow inhibition to include three layers: a bottom layer of independent Poisson processes modeling thalamus, and middle and top layers of integrate-and-fire neurons modeling V1 and MT, respectively (Fig. 3a and see Methods). We follow our past work with simplified firing rate networks^7^ and model a top-down attentional signal as an overall static depolarization to inhibitory neurons in the MT layer (Fig. 3a). This mimics cholinergic pathways that primarily affect interneurons^35,36^ and are thought to be engaged during attention^7,13^. The increased recruitment of inhibition during attention reduces the population-wide fluctuations in the MT layer (Fig. 3b) and decreases pairwise spike count correlations of MT-MT neuron pairs (Fig. 3c), while simultaneously increasing the correlation of V1-MT neuron pairs (Fig. 3d). Thus, this simple implementation of attentional modulation^7^ nonetheless captures the main aspects of the V1-MT dataset (Fig. 1d;^23^). Further, neuron pairs with larger firing rate increases also show larger correlation reductions (Fig. S3), in agreement with population recordings during both spatial and feature attention^37^.

**Figure 3:**
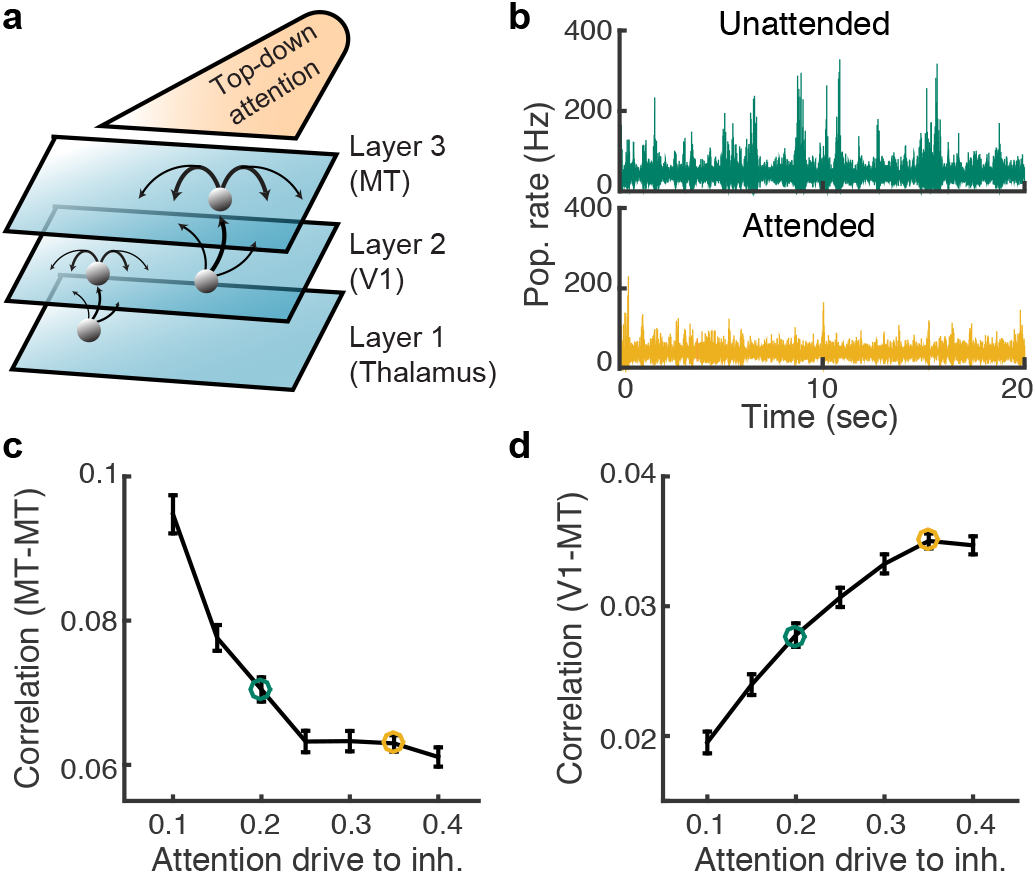
Top-down depolarization of MT inhibitory neurons capture the differential attentional modulation of shared variability within and across V1 and MT. **a**, Thalamus, V1, and MT are modeled in a three layer hierarchy of spatially ordered balanced networks. Top-down attentional modulation is modeled as a depolarization to MT inhibitory neurons (*μ_I_*). In both V1 and MT the recurrent projections are broader than feedforward projections and recurrent inhibition is slower than excitation. **b**, Population averaged firing rate fluctuations from MT in the unattended state (*μ_I_* = 0.2, green) and the attended state (*μ_I_* = 0.35, orange). **c**, Mean spike count correlation (*r_SC_*) of excitatory neuron pairs in MT decreases with attentional modulation. **d**, Mean *r_SC_* between the excitatory neurons in MT and the excitatory neurons in V1 increases with attention. Error bars are SEM.

Before we expose the core mechanisms through which attention modulates correlated activity we first give a broader analysis of shared variability in both our data and model. To this end we use dimensionality reduction tools^8,38^ to study the population-wide structure of trial-to-trial variability, rather than focusing only on individual pairwise correlation coefficients. We partition the covariance matrix into the shared variability among the population and the private noise to each neuron; the eigenvalues of the shared covariance matrix represent the variance along each dimension (or latent variable), while the corresponding eigenvectors represent the projection weights of the latent variables onto each neuron (see Methods). Applying these techniques to the multi-electrode V4 data^24^ shows a single dominant eigenmode (Fig. 4a, top, single session result see Fig. S4). This mode influences most of the neurons in the population in the same way (Fig. 4a middle, weights are dominant positive), and after subtracting the first mode the mean residual covariances are very small (Fig. 4a bottom). Finally, attention affects population variability primarily by quenching this dominant mode (Fig. 4a top, orange vs green) and the attentional modulation in the dominant mode is highly correlated with the modulation in mean covariance (Fig. S4c). The low dimensional structure of shared variability in our data is consistent with similar analysis in other cortices^4,6,8^, as well as alternative analysis of the same V4 data using generalized point process models^5^.

**Figure 4:**
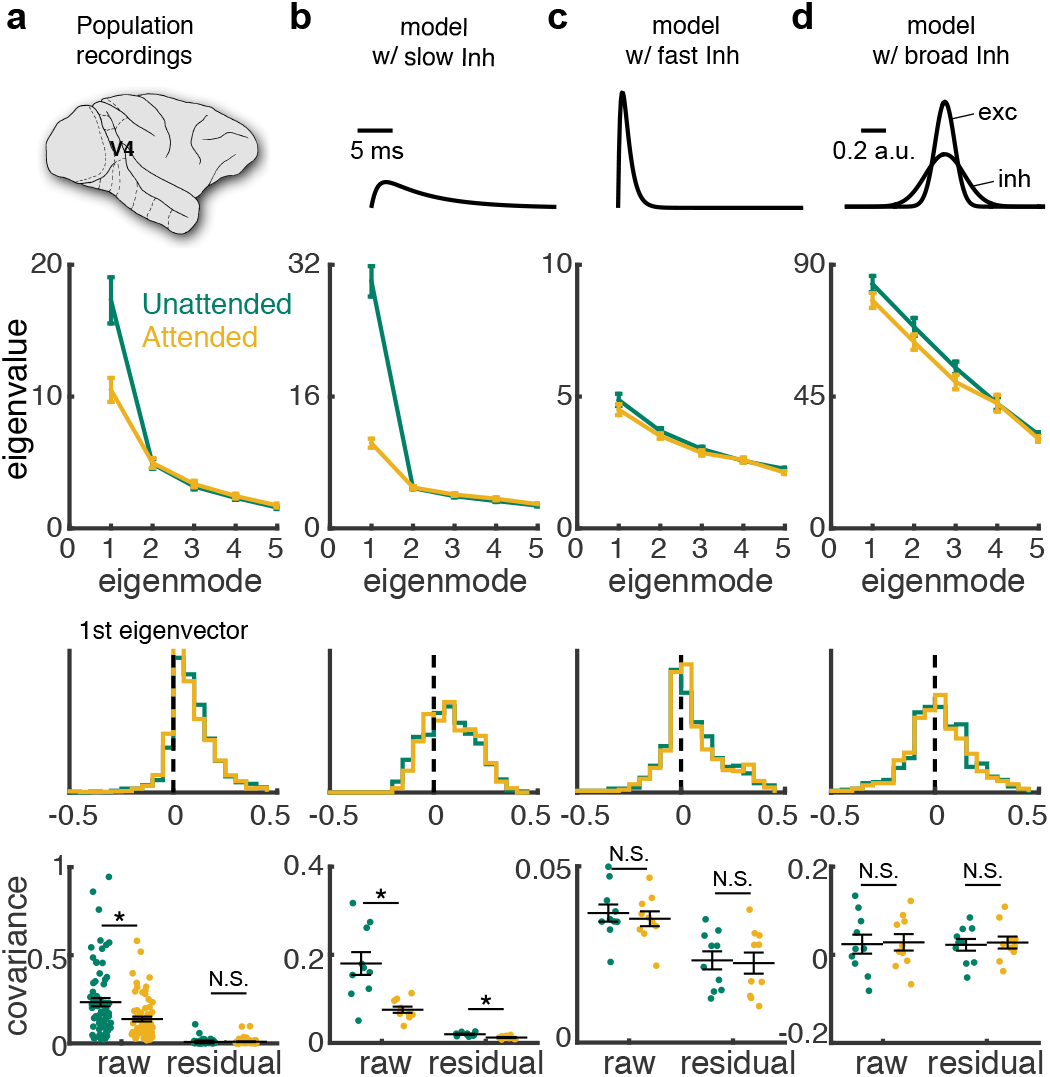
Internally generated shared variability from the model network is low-dimensional. **a**, Top: The first five largest eigenvalues of the shared component of the spike count covariance matrix from the V4 data^24^. Green: unattended; orange: attended; data from n=72 sessions with 43 ± 15 neurons. Error bars are SEM. Middle: the vector elements for the first (dominant) eigenmode. Bottom: the mean covariance from each session in attended and unattended states before (raw) and after (residual) subtracting the first eigenmode (mean ± s.e.m. in black). **b-d**, Same as **a** but for the three layer model with slow inhibition (**b**), model with fast inhibition (**c**) and model with slow and broad inhibition (**d**); n=10 samples of 50 neurons each. Two-sided Wilcoxon rank-sum test (attended vs unattended): mean covariance, **a**, *P* = 0.0013, **b**, *P* = 1.78 × 10^−22^, **c**, *P* = 0.7798 and **d**, *P* = 0.5850; residual, **a**, *P* = 0.7477, **b**, *P* = 5.40 × 10^−4^, **c**, *P* = 0.8796 and **d**, *P* = 0.5326.

The dimensionality of shared variability offers a strong test for our cortical model. We analyzed the spike count covariance matrix constructed from a subsampling of the spike trains in the third layer of our network model (*n* = 50 neurons). The network with slow inhibition produced shared variability with a clear dominant eigenmode that mimicked many of the core features observed in the V4 data (Fig. 4b). Further, the top-down attentional modulation of inhibition also suppressed this dominant mode (Fig. 4b top, orange vs green). The agreement between model and data broke down when inhibitory kinetics were faster than those of excitation, as was the case in our past studies^25,26,29^. Here, shared variability did not have a dominant mode (Fig. 4c, top), the raw mean correlation coefficient was near zero (Fig. 4c, bottom), and attentional modulation had a negligible effect on population variability (Fig. 4c, orange vs green). Experimental measurements of local cortical circuitry show that excitation and inhibition project on similar spatial scales^31,33^. When the model inhibitory projections in the third layer were spatially broader than those of excitation, thus at odds with experiment, then the model again disagreed with our V4 data (Fig. 4d). In sum, the low dimensional structure of shared variability requires inhibition that is neither faster nor anatomically broader than excitation – both features of real cortical circuits^28,31,39^. Further, a simple recruitment of inhibition through top-down drive can restore stability and quench low dimensional population variability.

This success of our model is quite distinct from that of past studies where low dimensional correlated variability was imposed from outside sources^3,7,19–21^. Rather, the shared variability in our model is internally generated from recurrent network interactions. We next explore how the inherent nonlinear dynamics that produce this variability allow our model to satisfy the constraints imposed by the differential correlation modulation of the within area and between area pairs (Fig. 1e).

### Relating low dimensional variability to spatio-temporal pattern formation

Networks of spiking neuron models produce rich activity that can be directly compared to population recordings. However, when these networks are outside the asynchronous regime they are not easily amenable to a deeper mechanistic analysis. An often used simplification to spiking dynamics are firing rate models where network interactions are mediated only through dynamic firing rates^1,40^. While these models lack a principled connection to spiking network models, they do produce qualitatively similar dynamics in recurrent networks and their simplicity makes them amenable to analysis techniques from dynamical systems theory. To gain intuition about how recurrent circuitry shapes low dimensional shared variability we considered a firing rate model that incorporated both the spatial architecture and synaptic dynamics that were central to our spiking model (see Methods).

Solutions where firing rates are constant over time are interpreted as asynchrony within the network, since only dynamical co-fluctuations in firing rates would mimic correlated spiking. We focused on how the stability of the asynchronous firing rate solution depended upon the temporal (*τ_i_*) and spatial (*σ_i_*) scales of inhibition. A firing rate solution is stable if the linearized dynamics are such that every eigenmode has eigenvalues with strictly negative real part. Since our network is spatially ordered the eignemodes are also organized in space, each with their own distinct spatial frequency. If the solution loses stability at a particular eigenmode, then the spatio-temporal dynamics of the resulting network firing rates will inherit the spatial frequency of that eigenmode – this process is termed spatio-temporal pattern formation^41^.

If *τ_i_* and *σ_i_* are near those of recurrent excitation, then a stable firing rate solution exists (Fig. 5a, grey region; 5b, top left, black curve with *τ_i_* = 5ms). Our past work explored activity within this regime^26^. When *τ_i_* increases and excitation and inhibition project with the same spatial scale (*σ_i_* = *σ_e_*), firing rate stability is first lost at an eigenmode with zero spatial frequency (Fig. 5b, top left). This creates population dynamics with a broad spatial pattern, allowing variability to be shared over the entire network. Simulations of the three layered spiking network model in this regime shows turbulent dynamics that extend across the entire network (Fig. 5b, top right; Supplementary movie S3). The projection weights of the first eigenmode from factor analysis of the network of spiking neuron models (Fig. 4b) show a uniform distribution in space (Fig. 5b, bottom), consistent with shared fluctuations of low spatial frequency. In contrast to this case, when *τ_i_* increases yet inhibition projects lateral to excitation (*σ_i_* > *σ_e_*), stability is first lost at a nonzero spatial frequency (Fig. 5c, top left). This creates population dynamics with coherence over a band of higher spatial frequencies, producing higher dimensional shared variability, as evident in the spatially patchy turbulent spiking dynamics of the three layered spiking network in this regime (Fig. 5c, top right; Supplementary movie S4). Correspondingly, the projection weights of the first three eigenmodes (Fig. 4d) show patterns of higher spatial frequency (Fig. 5c, bottom). Thus, the spatial and temporal scales of inhibition determine in large part the spatio-temporal patterns of network activity. Further, we now understand from a theoretical viewpoint why slow inhibition that does not project lateral to excitation is needed to account for the spiking data in both the network of spiking neuron models and experiment.

**Figure 5:**
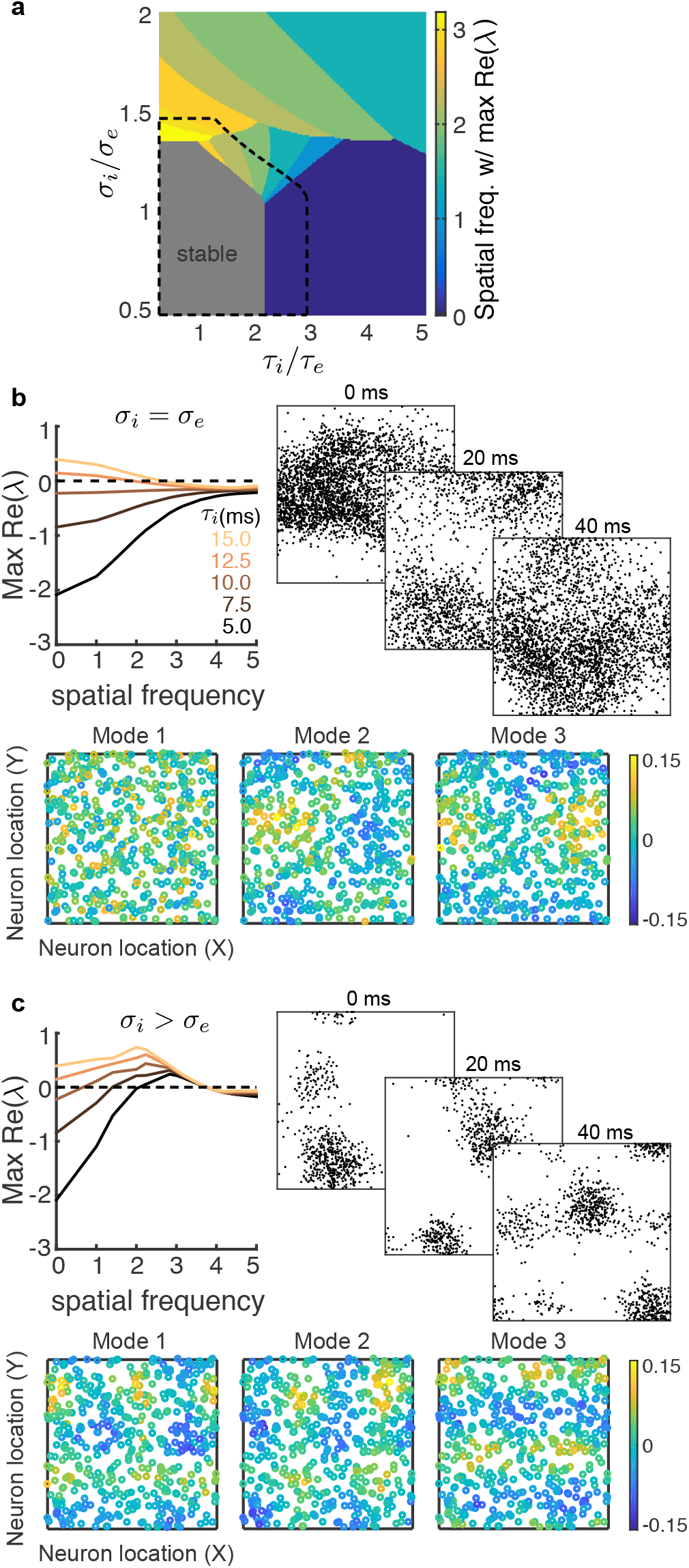
Stability analysis of a two-dimensional firing rate model. **a**, Bifurcation diagram of a firing rate model as a function of the inhibitory decay time scale *τ_i_* and inhibitory projection width *σ_i_*. The excitatory projection width and time constant are fixed at *σ_e_* = 0.1 and *τ_e_* = 5 ms, respectively. Color represents the spatial frequency with the largest real part of eigenvalue and the gray region is stable. Top-down modulation of inhibitory neurons modeling attention expands the stable region (black dashed). **b**, Top left: the real part of eigenvalues as a function of spatial frequency for increasing *τ_i_* when *σ_i_* = *σ_e_*. Top right: three consecutive spike raster snapshots of a spiking neuron network with *σ_i_* = *σ_e_* and slow inhibition (same network as in Fig. 4b in the unattended state). Bottom: spatial structure of projection weights from the first three eigenmode from Factor analysis of the spiking neuron network as in top right (n=500 neurons). **c**, Same as **b** for *σ_i_* larger than *σ_e_*. Top right and Bottom: same network as in Fig. 4d in the unattended state.

Finally, in the firing rate network we can also model attention as a depolarization to the inhibitory neurons, as was done in the network of spiking neuron models. In the firing rate network, attentional modulation expanded the stable region in the bifurcation diagram (Fig. 5a, dashed black line). In other words, attention increased the domain of firing rate stability. Thus, with *τ_i_* > *τ_e_* chosen so that in the unattended state the network was unstable at a low spatial frequency yet with attention the network was in the stable regime, our model captures the large attention-mediated quenching of population-wide shared variability reported in the population recordings (Fig. 4a) and network of spiking neuron models (Fig. 4b).

### Chaotic population-wide dynamics reflects internally generated variability

The attention-mediated differential modulation of within and between area correlations lead us to propose that shared variability has a sizable internally generated component. Using our heuristic model we argued that attention must quench a significant component of the variability (Var(*H*)) to account for the population recordings (Fig. 1e, bottom). This is a difficult constraint to satisfy and requires the mechanisms that produce internally generated variability to sensitively depend on top-down modulations. The firing rate model captured this sensitivity through a spatio-temporal pattern forming transition in network activity. However, the firing rate model does not internally produce trial-to-trial variability that can be compared to experiment, and we thus return to analysis of the network of spiking neuron models to probe how trial-to-trial variability is internally generated through recurrent coupling.

To isolate the sources of externally and internally generated fluctuations in the third layer of our network we fixed the spike train realizations from the first layer (thalamic) neurons as well as the membrane potential states of the second layer (V1) neurons, and only the initial membrane potentials of the third layer (MT) neurons were randomized across trials (Fig. 6a). This produced deterministic network dynamics when conditioned on activity from the first two layers, and consequently any trial-to-trial variability is due to mechanics internal to the third layer.

**Figure 6:**
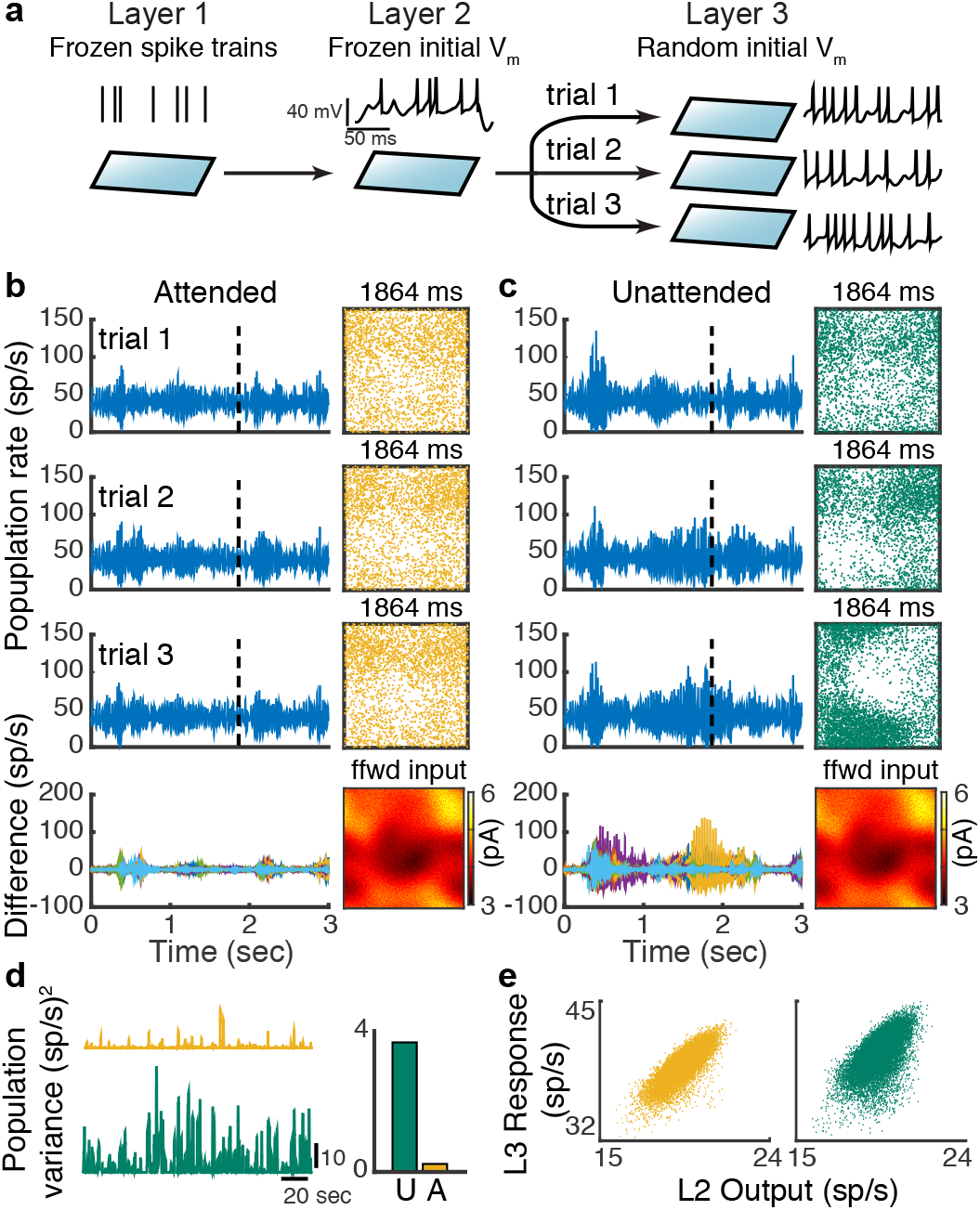
Chaotic population firing rate dynamics is quenched by attention. **a**, Schematic of the numerical experiment. The spike train realizations in layer one and the initial states of the membrane potential of layer two neurons are identical across trials, while in each trial we randomized the initial states of the layer three neuron’s membrane potentials. **b**, Three representative trials of the layer three excitatory population rates in the attended state (left row 1-3). Bottom row: difference of the population rates across 20 trials. Right (row 1-3): Snapshots of the neuron activity at time point 1864 ms. Each dot is a spike within 2 ms window from the neuron at that location. Right bottom: the synaptic current each layer three neuron receives from layer two at time 1864 ms. **c**, Same as **b** for the network in the unattended state. **d**, Trial-to-trial variance of layer three population rates as a function of time; right: mean variance across time. **e**, The layer three population rate tracks the layer two population rate better in the attended state. Both outputs and responses are smoothed with a 200 ms window.

The spike trains from third layer neurons in both the unattended and attended states have significant trial-to-trial variability despite the frozen layer one and two inputs. This is reflective of a well studied chaotic network dynamic in balanced networks where the spike times from individual neurons are very sensitive to perturbations that affect the spiking of other neurons^27,42^. To investigate how this microscopic (single neuron) variability possibly manifests as macroscopic population activity, we considered the trial-to-trial variability of the population-averaged instantaneous firing rate. While the population firing rate is dynamic in the attended state, there is very little variability from trial-to-trial (Fig. 6b, left; Fig. 6d, orange). A consequence of this low population-wide variability is the faithful tracking of the spatiotemporal structure of layer two outputs by layer three responses (Fig. 6b, right; e, orange). This tracking reflects the higher correlation between layer two and three spiking in the attended state (Fig. 3c). In contrast, in the unattended state the asynchronous solution is unstable, resulting in population-wide recruited activity. These periods of spatial coherence across the network are not trial locked and rather contribute to sizable trial-to-trial variability of population activity (Fig. 6c, left; d, green). This degrades the tracking of layer two outputs (Fig. 6c, right; e, green) and ultimately lowers the correlation between layer two and three spiking (Fig. 3c). Taken together, while the network model is chaotic in both the attended and unattended states, the chaos is population-wide only in the inhibition deprived unattended state.

The nonlinear pattern forming dynamics of the spatially distributed recurrent network impart extreme sensitivity to the population-wide internally generated variability. Indeed, in our model the trial-to-trial population rate variability is almost extinguished with attention (Fig. 6d, right). In our heuristic model with hidden variable *H* this amounts to Var(*H*) reducing drastically with attention, which is precisely what is needed to account for the differential modulation of within and between area correlations (Fig. 1e).

## Discussion

There is a longstanding research program aimed at understanding how variability is an emergent property of recurrent networks^16,17,25–27,42^. However, models are often restricted to simple networks with disordered connectivity. Consequently, population-wide activity is asynchronous, at odds with many experimental findings^2,3^. A parallel stream of research focuses on spatiotemporal pattern formation in neuronal populations, with a rich history in both theoretical^40^ and experimental contexts^43^. Yet a majority of these studies consider only trial-averaged activity, with tacit assumptions about how spiking variability emerges (but see^44^ and^26^). In this study we combined these modelling traditions with the goal of circuit-based understanding of the genesis and modulation of low dimensional internally generated shared cortical variability.

### Population-wide variability in balanced networks

Our model extends classical work in balanced cortical networks^16,18^ to include two well accepted experimental observations. First, cortical connectivity has a wiring rule that depends upon the distance between neuron pairs^31,32^. Theoretical studies that model distance dependent coupling commonly assume that inhibition projects more broadly than excitation^40,44,45^ (but see^34^). However, measurements of local cortical circuitry show that excitation and inhibition project on similar spatial scales^31,33^, and long-range excitation is known to project more broadly than inhibition^46^. Our work shows that this architecture is required for internally generated population variability to be low dimensional (Fig. 4b, d). The second observation is that inhibition has temporal kinetics that are slower than excitation^28^. Past theoretical models of recurrent cortical circuits have assumed that inhibition is not slower than excitation^16,18,34,47^, including past work from our group^26^. Consequently, these studies could only capture the residual correlation structure of population recordings once the dominant eigenmode was subtracted^8,26^; in these cases the residual accounted for less than ten percent of the true shared variability. The asynchronous solution is unstable when inhibition is slower than excitation, and in networks with two spatial dimensions the resulting dynamics are weakly correlated, matching experiments (Fig. 2 and 4). In total, by including accepted features of cortical anatomy and physiology, long ignored by theorists, our model network recapitulates low dimensional population-wide variability to a much larger extent than previous models.

The above narrative is somewhat revisionist; there are several well known theoretical studies in disordered networks where one dimensional population-wide correlations do emerge, notably in networks where rhythmic^17^ or ‘up-down’^45,47^ dynamics are prominent. Networks with dense yet disordered connectivity ensure that all neuron pairs receive some shared inputs from overlapping presynaptic projections. In such a network if the asynchronous state becomes unstable then this shared wiring will correlate spiking activity across the entire network. In other words, any shared variability will be one dimensional (scalar) by construction. In contrast, the ordered connectivity in our network is such that neuron pairs that are distant from one another have no directly shared presynaptic connections. Consequently, when asynchrony is unstable one dimensional population dynamics is not preordained, rather the spatial network can support higher dimensional shared variability depending on the temporal and spatial scales of recurrent coupling (Fig. 4b,c; Fig. 5). From the vantage of this model we discovered the conditions for recurrent architecture and synaptic physiology for low dimensional shared variability

### Internal versus external population variability

Our circuit model assumed that the component of population-wide variability that is subject to attentional modulation was internally generated within the network. This was motivated by constraints imposed by the differential attentional modulation of within and between pairwise correlations in our population recordings^23^ (Fig. 1). While our model is a parsimonious explanation of the data, it does not definitively exclude mechanisms where variability is inherited from outside sources. In fact it is difficult to conceive of descending synaptic and cholinergic projections from higher areas that would not contribute some trial-to-trial variability to a receiving neuronal population.

Fluctuations from external sources are an often assumed and straightforward mechanism for population-wide variability^3,7,19–21,48^. However, if this framework aims to capture a modulation in variability a further choice must be made^3^. One way to modulate population-wide variability is to simply allow the amplitude of input fluctuations to change. Such an ‘inheritance model’ is often assumed for how top-down feedback to either visual areas V1^48^ or MT^21^ determines choice probability in ambiguous decision tasks. When V2 and V3 are inactivated through cooling the single neuron variability in MT is markedly reduced suggestive that a component of variability is feedforward propagated^49^. This is in contrast to the only slight reductions in V1 variability when feedback projections from V2 and V3 are inactivated^49^. Thus, there is limited experimental evidence for direct top-down contributions to single neuron variability. Additional multi-area population recordings between connected brain regions will be needed to probe how correlated variability flows along bottom-up and top-down pathways.

The second way to change population output variability is to keep input fluctuations fixed yet shift the operating point of the network so that the nonlinearties inherent in spiking dynamics change input-output transfer of variability. This mechanism has been suggested for how top-down attentional modulation affects population variability in recurrent excitatory-inhibitory cortical networks^7,19^. Network models with either disordered connectivity or simple one dimensional spatial structure must have a stable asynchronous state, else the internally generated correlations are excessive (Fig. 2aiii,c). Consequently, when such networks are used to model attentional modulation both the attended and unattended states must be in the asynchronous regime^7,19^. In such cases, population-wide variability must be from outside the network and attention only changes how the network filters these external fluctuations.

In contrast, the two dimensional spatial structure in our model supports rich chaotic network dynamics outside the asynchronous state, yet with population-wide correlations that are a reasonable mimic of experiment (Fig. 2aiv,c). Spatiotemporal chaos is a hallmark feature of systems that are far from equilibrium in physics, chemistry and biology^41^. In particular, low viscosity fluids produce a special brand of spatiotemporal chaotic behavior labelled turbulence, characterized by the presence of vortices and eddies in the fluid flow^50^. Like our network, the character of turbulent flow is very dependent upon the dimension of the fluid, with one dimensional fluids not showing turbulence, and two dimensional turbulent flow having larger spatial scales than the flow in full three dimensional fluids^50^. The dynamics within recurrent networks of neurons are certainly not equivalent to that of fluids, in part because they possess both short and long range interactions in contrast to only the direct local interactions in fluids. Nevertheless, the fluid analogy to our work is tempting since the chaotic dynamics of our two dimensional network has a macroscopic character that permits low, but non-vanishing, microscopic correlations, in contrast to the unrealistic high correlation dynamics of one dimensional or disordered networks. While the top-down attentional signal in our model is similar to that used in simpler models^7,19^, the effect of top-down attention is to not only shift the operating point of the network but also dampen the macroscopic chaotic dynamics of the network. In other words, attention not only attenuates the transfer of population-wide variability but also quenches the variability that is to be transferred. This permits a near complete attention-mediated suppression of internally generated correlations (Fig. 6). This extreme sensitivity allows top-down inputs to easily control the processing state of a network.

State dependent shifts in population-wide variability are widespread throughout cortex^3^, and are often a signature of cognitive control. The circuit structure of our network is not a special feature of the primate visual system, yet rather a generic property of most cortices. We thus expect that the basic mechanisms for population-wide variability and its modulation exposed in our study will be operative in many regions of the cortex, and in many animal systems.

## Methods

### Network model description

The network consists of three layers. Layer 1 is modeled by a population of *N*_1_ = 2, 500 excitatory neurons, the spikes of which are taken as independent Poisson processes with a uniform rate *r*_1_ = 10 Hz. Layer 2 and Layer 3 are recurrently coupled networks with excitatory (*α* = e) and inhibitory (*α* = i) populations of *N*_e_ = 40, 000 and *N*_i_ = 10,000 neurons, respectively. Each neuron is modeled as an exponential integrate-and-fire (EIF) neuron whose membrane potential is described by:

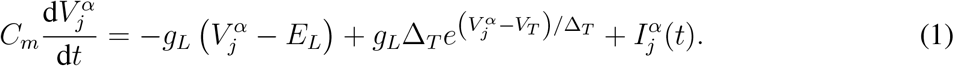

Each time 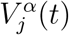 exceeds a threshold *V*_th_, the neuron spikes and the membrane potential is held for a refractory period *τ*_ref_ then reset to a fixed value *V*_re_. Neuron parameters for excitatory neurons are *τ_m_* = *C_m_*/*g_L_* = 15 ms, *E_L_* = −60 mV, *V_T_* = −50 mV, *V*_th_ = −10 mV, Δ_*T*_ = 2 mV, *V*_re_ = −65 mV and *τ*_ref_ = 1.5 ms. Inhibitory neurons are the same except *τ_m_* = 10 ms, Δ_*T*_ = 0.5 mV and *τ*_ref_ = 0.5 ms. The total current to each neuron is:

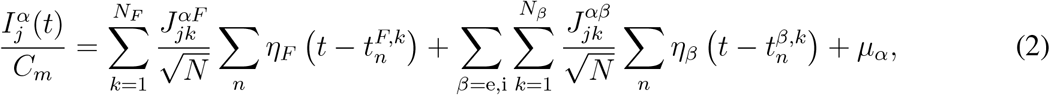

where *N* = *N_e_* + *N_i_* is the total number of the network population. Postsynaptic current is

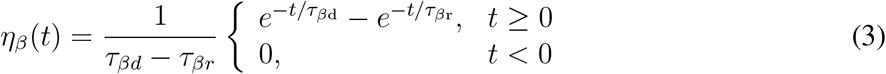

where *τ*_er_ = 1 ms, *τ*_ed_ = 5 ms and *τ*_ir_ = 1 ms, *τ*_id_ = 8 ms. The feedforward synapses from Layer 1 to Layer 2 have the same kinetics as the recurrent excitatory synapse, i.e. 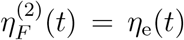. The feedforward synapses from Layer 2 to Layer 3 have a fast and a slow component.

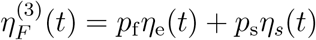

with *p*_f_ = 0.2, *p*_s_ = 0.8. *η_s_*(*t*) has the same form as Eq. 3 with a rise time constant 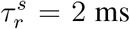 and a decay time constant 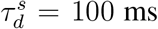. The excitatory and inhibitory neurons in Layer 3 receive static current *μ_e_* and *μ_i_*, respectively.

Neurons on the three layers are arranged on a uniform grid covering a unit square Γ = [0, 1] × [0, 1]. The probability that two neurons, with coordinates **x** = (*x*_1_, *x*_2_) and **y** = (*y*_1_, *y*_2_) respectively, are connected depends on their distance measured periodically on Γ:

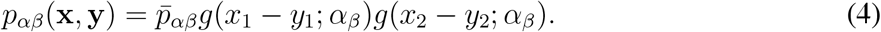

Here 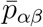 is the mean connection probability and

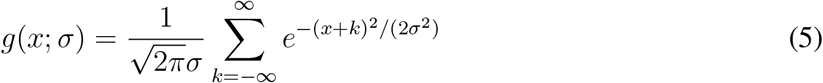

is a wrapped Gaussian distribution. Excitatory and inhibitory recurrent connection widths of Layer 2 are 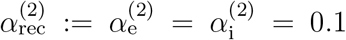 and feedforward connection width from Layer 1 to Layer 2 is 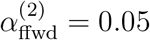. The recurrent connection width of Layer 3 is 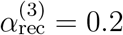 and the feedforward connection width from Layer 2 to Layer 3 is 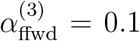. A presynaptic neuron is allowed to make more than one synaptic connection to a single postsynaptic neuron.

The recurrent connectivity of Layer 2 and Layer 3 have the same synaptic strengths and mean connection probabilities. The recurrent synaptic weights are *J*_ee_ = 80 mV, *J*_ei_ = −240 mV, *J*_ie_ = 40 mV and *J*_ii_ = −300 mV. Recall that individual synapses are scaled with 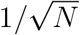 (Eq. 2); so that, for instance, 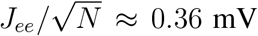. The mean connection probabilities are 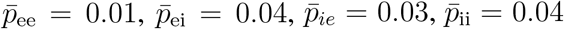. The out-degrees are 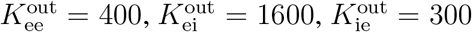 and 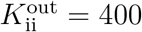. The feedforward connection strengths from Layer 1 to Layer 2 are 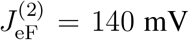 and 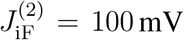 with probabilities 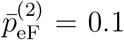 and 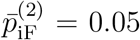 (out-degrees 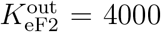 and 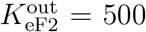). The feedforward connection strengths from Layer 2 to Layer 3 are 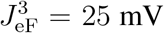 and 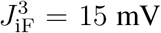 with mean probabilities 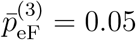 and 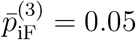 (out-degrees are 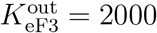 and 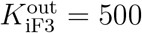). Only the excitatory neurons in Layer 2 project to Layer 3.

The spatial models in Fig. 2aii,aiv contain only Layer 1 and Layer 2. In the model with disordered connectivity, the connection probability between a pair of neurons is 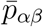, independent of distance. Other parameters are the same as the spatial model. The decay time constant of IPSC (*τ*_id_) was varied from 1 to 15 ms (Fig. 2d). The rise time constant of IPSC (*τ*_ir_) is 1 ms when *τ*_id_ > 1 ms and 0.5 ms when *τ*_id_ = 1 ms.

The parameters used in Fig. 3c,d are *μ_i_* = [0.1, 0.15, 0.2, 0.25,0.3, 0.35, 0.4] pA and *μ_E_* = 0 pA. The mean firing rates in Layer 2 are 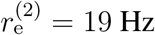 and 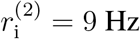. In the further analysis (Fig. 4b-d, Fig. 5b and Fig. 6), we used *μ_I_* = 0.2 pA for the unattended state and *μ_I_* = 0.35 pA for the attended state. In simulations of the spatial model with fast inhibition (Fig. 4c), *τ*_ir_ = 0.5 ms, *τ*_id_ = 1 ms. In simulations of the spatial model with broad inhibitory projection (Fig. 4d and Fig. 5c), 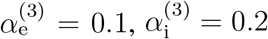. Other parameters are not changed.

All simulations were performed on the CNBC Cluster in the University of Pittsburgh. All simulations were written in a combination of C and Matlab (Matlab R 2015a, Mathworks). The differential equations of the neuron model were solved using forward Euler method with time step 0.01 ms.

### Neural field model and stability analysis

We use a two dimensional neural field model to describe the dynamics of population rate (Fig. 5). The neural field equations are

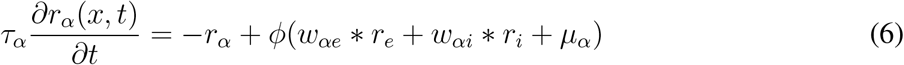

where *r_α_*(*x, t*) is the firing rate of neurons in population *α* = e, i near spatial coordinates *x* ∈ [0, 1] × [0, 1]. The symbol ∗ denotes convolution in space, *μ_α_* is a constant external input and 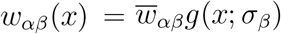 where *g*(*x*; *σ_β_*) is a two-dimensional wrapped Gaussian with width parameter *σ_β_, β* = e, i. The transfer function is a threshold-quadratic function, *ϕ*(*x*) = [*x*^2^]_+_. The timescale of synaptic and firing rate responses are implicitly combined into *τ_α_*. In networks with approximate excitatory-inhibitory balance, rates closely track synaptic currents^18^, so *τ_α_* represents the synaptic time constant of population *α* = e, i.

For constant inputs, *μ_e_* and *μ_i_*, there exists a spatially uniform fixed point, which was computed numerically using an iterative scheme^25^. Linearizing around this fixed point in Fourier domain gives a Jacobian matrix at each spatial Fourier mode^25^

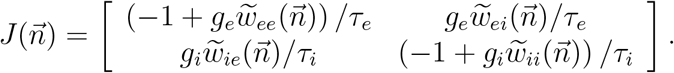

where 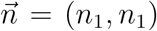 is the two-dimensional Fourier mode, 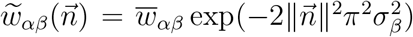 is the Fourier coefficient of *w_αβ_*(*x*) with 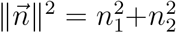 and *g_a_* is the gain, which is equal to *ϕ*′(*r_α_*) evaluated at the fixed point. The fixed point is stable at Fourier mode 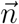 if both eigenvalues of 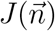 have negative real part. Note that stability only depends on the wave number, 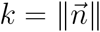, so Turing-Hopf instabilities always occur simultaneously at all Fourier modes with the same wave number (spatial frequency).

For the stability analysis in Fig. 5a, *τ_i_* varies from 2.5 ms to 25 ms, *σ_i_* varies from 0.05 to 0.2, and *τ_e_* = 5 ms and *σ_e_* = 0.1. The rest of the parameters were 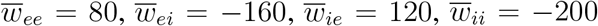, *μ_e_* = 0.48 and *μ_i_* = 0.32. Depolarizing the inhibitory population (*μ_I_* = 0.5) expands the stable region (Fig. 5a, black dashed).

### Experimental methods

Each of the two datasets (recordings from V4 and recordings from V1 and MT) was collected from two different rhesus monkeys as they performed an orientation-change detection task. All animal procedures were in accordance with the Institutional Animal Care and Use Committee of Harvard Medical School, University of Pittsburgh and Carnegie Mellon University.

For analysis in Fig. 1b and Fig. 4a, data was collected with two microelectrode arrays implanted bilaterally in area V4^24^. In our analysis, we include stimulus presentations prior to the change stimulus from correct trials, excluding the first stimulus in a trial to avoid adaptation effects. Spike counts during the sustained response (120 - 260 ms after stimulus onset) are considered for the correlation and factor analysis. Neurons recorded from either the left or right hemisphere in one session are treated separately. There are a total of 42,496 trials for 72,765 pairs from 74 recording sessions. Two sessions from the original study were excluded for factor analysis due to inadequate trials. The trial number and unit number of each session is summarized in Table S1.

For analysis in Fig. 1d, data was collected with one microelectrode array implanted in area V1 and a single electrode or a 24-channel linear probe inserted into MT^23^. Again, our analysis includes full contrast stimulus presentations prior to the change stimulus from correct trials and excludes the first stimulus in a trial to avoid adaptation effects. Spike counts are measured 30 - 230 ms after stimulus onset for V1 and 50 - 250 ms after stimulus onset for MT to account for the average visual latencies of neurons in both areas. There are a total of 1,631 V1-MT pairs from 32 recording sessions.

### Statistical methods

To compute the noise correlation of each simulation, 500 neurons were randomly sampled without replacement from the excitatory population of Layer 3 and Layer 2 within a [0, 0.5]x[0, 0.5] square (considering periodic boundary condition). Spike counts were computed using a sliding window of 200 ms with 1 ms step size and the Pearson correlation coefficients were computed between all pairs. Neurons of firing rates less than 2 Hz were excluded from the computation of correlations. In Fig. 3c,d, for each *μ_i_* there were 50 simulations and each simulation was 20 sec long. Connectivity matrices and the initial states of each neuron’s membrane potential were randomized in each simulation. The first 1 second of each simulation was excluded from the correlation analysis. Standard error was computed based on the mean correlations of each simulation. For simulations of Fig. 2d, there was one simulation of 20 seconds per *τ*_id_ and the connectivity matrices were randomized for each simulation. To compute the noise correlation, 1000 neurons were randomly sampled without replacement in the excitatory population of Layer 2 within a [0, 0.5]x[0, 0.5] square. Correlations are computed between firing rates that are smoothed with a Gaussian window of width 10 ms.

Factor analysis assumes spike counts of n simultaneously recorded neurons *x* ∈ ℝ^*n*×1^ is a multivariable Gaussian process

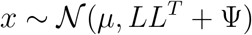

where *μ* ∈ ℝ^*n*×1^ is the mean spike counts, *L* ∈ ℝ^*n*×*m*^ is the loading matrix of the *m* latent variables and Ψ ∈ ℝ^*n*×1^ is a diagonal matrix of independent variances for each neuron. We choose *m* = 5 and compute the eigenvalues of *LL^T^, λ_i_* (*i* = 1, 2, …, 5), ranked in descending order. We compute the residual covariance after subtracting the first mode as

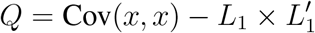

where Cov(*x, x*) is the raw covariance matrix of *x* and *L*_1_ is the loading matrix when fitting with *m* =1. The mean raw covariance and residual (Fig. 4a-d, bottom) are the mean of the off-diagonal elements of Cov(*x, x*) and *Q*, respectively. When applying factor analysis on model simulations (Fig. 4b-d), we randomly selected 50 excitatory neurons from Layer 3, whose firing rates were larger than 2 Hz in both the unattended and attended states. There were 10 non-overlapping sampling of neurons and we applied factor analysis on each sampling of neuron spike counts. There were 15 simulations with fixed connectivity matrices, each of which was 20 seconds long. Spike trains were truncated into 140 ms spike count window with a total of 2,025 counts per neuron. In simulations with fast inhibition (Fig. 4c) and broad inhibitory projection (Fig. 4d), the feedforward connectivity from Layer 2 to Layer 3 was the same as the one in simulations of the original model (Fig. 4b).

To study the chaotic population firing rate dynamics of Layer 3 (Fig. 6), we fixed the spike trains realizations from Layer 1 neurons, the membrane potential states of the Layer 2 neurons and all connectivity matrices. Only the initial membrane potentials of Layer 3 neurons were randomized across trials. There were 10 realizations of Layer 1 and Layer 2, each of which was 20 sec long. For each simulation of Layer 2, 20 repetitions with different initial conditions were simulated for Layer 3. The connectivity matrices in Layer 3 were the same across the 20 repetitions but different for each realization of Layer 1 and Layer 2. The realizations of Layer 1 and Layer 2 and the connectivity matrices were the same for the attended and unattended states. Trial-to-trial variance of Layer 3 population rates (Fig. 6d) was the variance of the mean population rates of the Layer 3 excitatory population, smoothed by a 200 ms rectangular filter, across the 20 repetitions. The first second of each simulation was discarded.

### Code availability

Computer code for all simulations and analysis of the resulting data is included in Supplementary Software.

### Data availability

The data that support the findings of this study are available from the corresponding author upon request.

## Supplemental Information

Figures. S1-S4

Table. S1

Supplementary Methods

Captions for Movies S1-S4

## Acknowledgments

NIH Grants CRCNS R01DC015139-01ZRG1 (B.D.), 4R00EY020844-03 (M.R.C.), R01 EY022930 (M.R.C.), 5T32NS7391-14 (D.A.R.), and Core Grant P30 EY008098; NSF Grants DMS-1517828, DMS-1654268 and DBI-1707400 (R.R.) and DMS-1517082 (B.D.); Vannevar Bush faculty fellowship N00014-18-1-2002 (B.D.), a Whitehall Foundation Grant (M.R.C.); a Klingenstein-Simons Fellowship (M.R.C.); grants from the Simons Foundation (B.D. and M.R.C.); a Sloan Research Fellowship (M.R.C.); a McKnight Scholar Award (M.R.C.).

## Author Contributions

C.H., M.R.C., and B.D. conceived the project; C.H. performed the simulations and data analysis; R.P. and R.R analyzed the firing rate model; D.A.R. and M.R.C. provided the experimental data; B.D. supervised the project; all authors contributed to writing the manuscript.

## Author Information

The authors declare no competing financial interests. Correspondence and requests for materials should be addressed to B.D.

